# Uncovering One-Dimensional Reaction Coordinate that Underlies Structure-Function Relationship of Proteins

**DOI:** 10.1101/2022.01.08.475519

**Authors:** Shanshan Wu, Huiyu Li, Ao Ma

**Author notes:** correspondence should be addressed to: Ao Ma.

## Abstract

Understanding the mechanism of functional protein dynamics is critical to understanding protein functions. Reaction coordinates is a central topic in protein dynamics and the grail is to find the one-dimensional reaction coordinate that can fully determine the value of committor (i.e. the reaction probability in configuration space) for any protein configuration. We present a powerful new method that can, for the first time, identify the rigorous one-dimensional reaction coordinate in complex molecules. This one-dimensional reaction coordinate is determined by a fundamental mechanical operator--the generalized work functional. This method only requires modest computational cost and can be readily applied to large molecules. Most importantly, the generalized work functional is the physical origin of the collectivity in functional protein dynamics and provides a tentative roadmap that connects the structure of a protein to its function.

---

Proteins are the building blocks of biological systems responsible for most biological functions. Understanding the mechanism of protein function is of paramount importance. The central dogma of protein science is that structure determines function. The holy grail is the physical principle that explains how the structure of a protein determines its function. The route to this principle starts with recognizing that most protein functions, such as ligand binding, allostery and effects of mutations, involve significant conformational dynamics because a protein has multiple functional structures and transitions between these structures are required for its function [1, 2]. Structure determines function because the specific structure enables the desired functional dynamics. Understanding protein function requires understanding functional dynamics.

## Importance of reaction coordinates

Most functional protein dynamics are activated processes similar to chemical reactions: a protein must cross an activation barrier much higher than thermal energy to move from the reactant (initial) state to the product (final) state. A central concept here is the reaction coordinates (**RC**s): the small number of essential coordinates that fully determine the progress of a reaction [3]. In particular, RCs determine the location of the transition state (i.e. the top of the activation barrier). The reaction progresses in the forward direction when RCs move towards the product state; it regresses when RCs move towards the reactant state; movements of all the other coordinates are irrelevant.

Reaction coordinates are the cornerstone of the standard reaction rate theories. Reaction dynamics is the dynamics of RCs. Indeed, Kramers theory is built on the assumption that the dynamics of the RC is governed by a Langevin equation [4], which is extended to generalized Langevin equation in Grote-Hynes theory [5]. Similarly, transition state theory relies on simplifying assumptions on the essential features of the dynamics of the RC [6–8]. The applicability of all these theories requires full knowledge of the RCs.

Reaction coordinates are also central to enhanced sampling, a major current research direction in computational biophysics. Most algorithms rely on applying biasing potentials on one or a few collective variables (**CV**s) selected by intuitions to increase sampling of the relevant configuration space [9–15]. However, these algorithms are effective only if the CVs coincide with RCs. Otherwise, the infamous “hidden barrier” problem will inevitably appear and prevent effective sampling [16–18]. The “hidden barrier” is the actual activation barrier that lies on the true RCs, which is missed by the biasing potentials along CVs that do not overlap with the RCs significantly.

## The rigorous concept of RC

Given the central importance of RCs, a rigorous definition is required. An important concept is the committor (*p_B_*) of a system configuration, defined as the probability that a dynamic trajectory initiated from this configuration, with initial momenta drawn from Boltzmann distribution, to reach the product state [3, 19–21]. Committor rigorously parameterizes the progress of a reaction (e.g. the transition state corresponds to *p_B_* = 0.5). Consequently, RCs are defined as the set of coordinates that fully determine the committor value of any system configuration. Other coordinates in the system are irrelevant. Reaction coordinates under this definition provide an accurate reduced description of a reaction. In contrast, ad hoc RCs that do not satisfy this definition bear little relevance to reaction dynamics and mechanism [22, 23].

A critical question is: How many RCs are there in a system? The standard reaction rate theories assume that, for a single-channel reaction, there exists a single one-dimensional RC (**1D-RC**) that can fully determine committor. Even though the multi-dimensional extensions of these theories assume that a reaction is controlled by the free energy surface of multiple RCs, it is always assumed that there is only one unstable direction in the saddle region, which is the direction of the 1D-RC. The seminal work by Berezhkovskii and Szabo [24] showed that it is possible to find a 1D-RC that can reproduce the same reaction rate determined by multiple RCs [25, 26].

In fact, the concept of multiple RCs is an approximation to the concept of 1D-RC. In the schematic example of Fig. 1a, *q*_1_ and *q*_2_ are considered RCs because they both have components along the direction of *r_c_*, the 1D-RC. However, they also have components that are orthogonal to *r_c_*, pointing to their inaccuracy as RCs. The concept of multiple RCs is semi-quantitative and has some leeway; the concept of 1D-RC is quantitative and precise. The former is adopted much more widely because, compared to the 1D-RC, they are much easier to identify.

**Figure 1:**
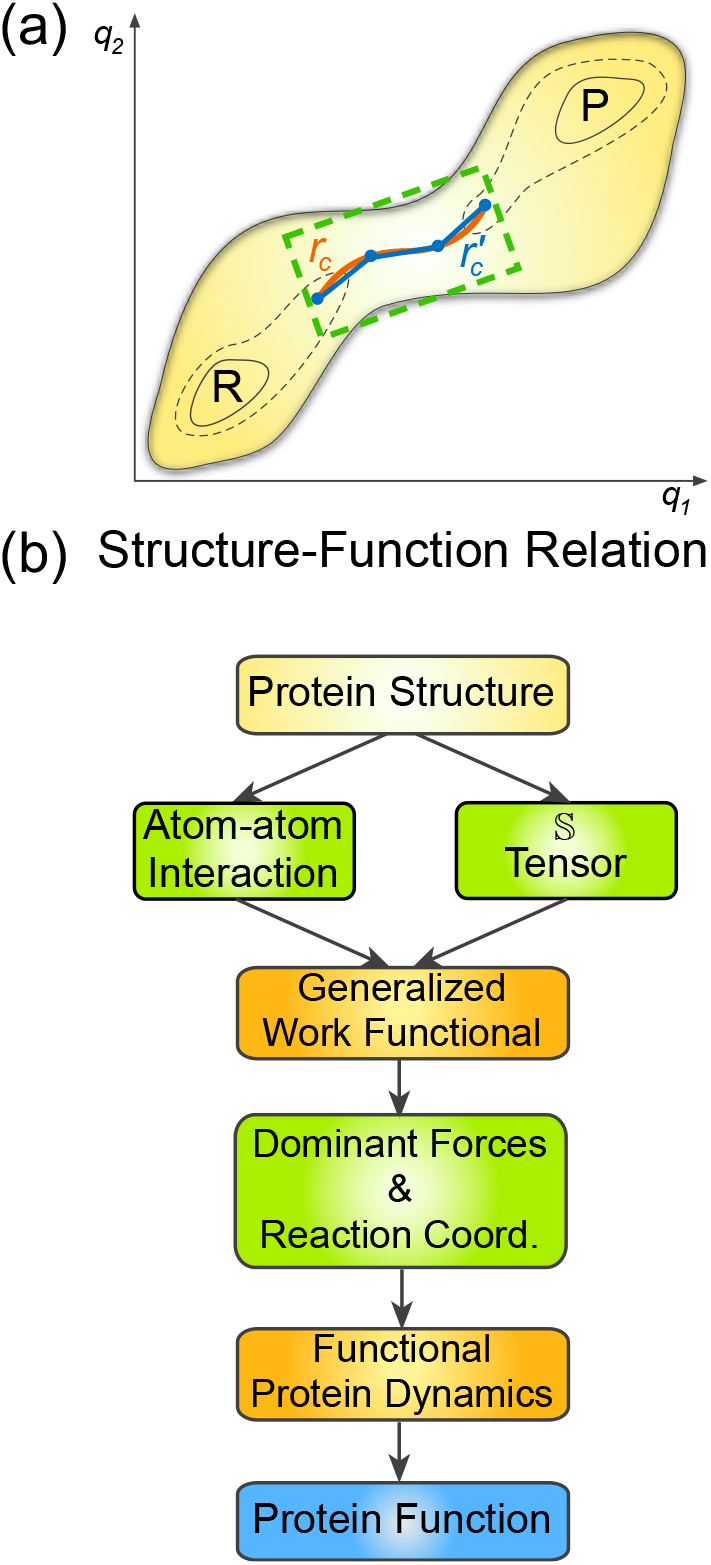
One-dimensional reaction coordinate and structure-funciton relationship. **(a**) A schematic of the potential energy surface and RCs of a two-dimensional system. The **gray** region indicates the contours of the potential energy surface, with the reactant and product states marked by **R** and **P** respectively. The **green** dashed rectangle marks the transition region. The **orange** curve indicates the 1D-RC and the **blue** line segments is its piecewise linear approximation. **(b)** Flow chart for the tentative roadmap for the physical principle that connects the structure of a protein with its function.

## The goal for studying RCs

The rigorously defined RCs provide a chance to bring order to the apparent chaos and complexity of protein dynamics that thwarted understanding of protein function. The small number of RCs compared to the myriad of coordinates in a protein indicates that protein dynamics has an intrinsic order and governing principles, which is embedded in the RCs as they accurately describe the progress of protein dynamics. Therefore, the physical cause behind the RCs is the physical determinant of this intrinsic order. The goal for studying RCs is to uncover and, furthermore, understand this physical determinant.

## Challenges in identifying RCs

The first step towards this goal is to identify the less restrictive multiple RCs, since it is easier than identifying the 1D-RC. However, it turned out to be a daunting challenge. Despite intensive efforts, over the past two decades there have been only a few successful cases where RCs that can determine committor were identified [27–30]. In each case, two or three RCs were successfully identified. The most important lesson from these examples is that RCs are often counter-intuitive [27, 28]. This counter-intuitiveness demonstrates why it is challenging to identify the few RCs out of practically infinitely many potential candidates in a complex molecule. For example, over 6,000 candidates were tested in ref. [28] for the *C_7eq_* → *α_R_* isomerization of an alanine dipeptide in solution to uncover the three RCs.

As a result, systematic methods that goes beyond intuition-based trial-and-error experimentation are required, as it is not feasible to enumerate the enormous number of potential candidates. The first systematic method used neural network and successfully identified the RCs for the *C_7eq_* → *α_R_* isomerization of an alanine dipeptide in solution that eluded human intuitions [28]. This early success is followed by many machine-learning methods along similar lines, a research direction that is currently attracting intensive attention but also facing significant challenges [23, 31–35]. More importantly, machine-learning methods cannot answer the central question concerning RCs: Why do RCs exist and how do they control protein dynamics? Answering this question requires the correct mechanical model of protein dynamics, which machine-learning cannot provide because it can only optimize parameters based on a pre-formulated model but cannot construct a model de novo. An approach based on fundamental physics is required.

## Physics-based approach for understanding RCs

The recently developed energy-flow theory offered some insights into the physical principle behind RCs [29, 36]. It showed that energy flows from the fast to the slow coordinates during a reaction [29]. Because RCs are the slowest, they carry the highest energy flows. This picture is intuitively appealing and explains why RCs are so important: energy is the currency of dynamics and movements of RCs incur the highest cost, thus they control reaction dynamics by dictating the overall cost. However, this picture is only semi-quantitative and does not answer a critical question in protein dynamics: What is the relationship between different RCs?

This question is automatically answered if we can identify the correct 1D-RC, because it is a function of the multiple RCs, precisely defining the relationship between them. Moreover, 1D-RC answers a key conceptual question: What is the physical origin of collectivity in functional dynamics? It is generally assumed that functional dynamics are controlled by one or a few CVs, which is also the reason that CVs are widely used in enhanced sampling. However, the critical questions have never been answered: Why functional dynamics are collective and what determine the CVs? By definition, 1D-RC is the optimal CV, thus the physical determinant behind 1D-RC is the physical origin of collectivity in protein dynamics.

In this paper, we introduced a fundamental mechanical operator rooted in Newton’s law, the generalized work functional (**GWF**), which determines the correct 1D-RC. Applying this new approach to a prototypical biomolecular isomerization process, the *C_7eq_* → *C_7ax_* transition of an alanine dipeptide in vacuum, we obtained, as an accurate approximation, the piecewise linearization of the 1D-RC (e.g. 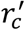 in the example of Fig. 1a) throughout the entire transition region (i.e. *p_B_* ∈ (0,1)). Each linear segment is a combination of four component RCs: two dihedrals and two improper dihedrals. This 1D-RC can predict the committor value throughout the entire range of *p_B_* with an accuracy that approaches the limit of the numerically evaluated committor values. This accuracy far exceeds what was achieved by previous methods. Among the four component RCs, the two improper dihedrals were only identified by the current method; their inclusion significantly improved the accuracy in predicting the committor value.

Most importantly, the GWF is a generic physical property universal to all protein molecules, allowing us to answer many fundamental questions concerning RCs and functional protein dynamics. 1) Why do RCs exist and how do they control protein dynamics? 2) What is the relationship between different RCs? 3) What is the origin of collectivity in protein dynamics?

The GWF provides an overall account of the mechanical effects of the couplings between different coordinates. It shows that, due to the special structural features of a protein, a small number of forces dominate the dynamics of the entire system. The directions of these forces are the RCs. Motions of RCs control these dominant forces, thus control the reaction dynamics. The GWF is a tensor with an inherent structure that can be revealed by singular value decomposition (**SVD**). The leading vector from SVD identifies the single dominant force; its direction is the 1D-RC, which is a function of all the RCs, thus coupling them with each other and synergizing their movements. This synergy between RCs is the origin of collectivity in functional protein dynamics because dynamics of RCs is the crux of protein dynamics.

From these results emerges a tentative roadmap that connects the structure of a protein to its function (Fig. 1b). Protein structure determines both atom-atom interactions and the structural coupling tensor responsible for the mechanical couplings between different coordinates. Together, these two factors determine the GWF, which determines the directions of the dominant forces that appear as the RCs. Finally, RCs control the functional dynamics, which determines function.

## Results

### Energy-Flow Theory

A major theoretical tool we use for analyzing reaction dynamics is the energy flow theory [29, 36]. This theory defines the potential energy flow through a coordinate *q_i_* as its work [29]:

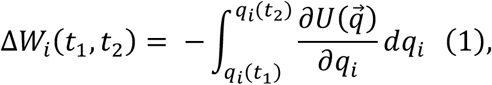

where 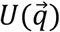 is the potential energy of the system, 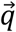 is the position vector of the system in the configuration space. According to Eq. (1), Δ*W_i_*(*t*_1_, *t*_2_) is the change in 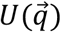 caused by the motion of *q_i_* alone along a dynamic trajectory in the time interval [*t*_1_, *t*_2_]. It is a projection of the change in the total potential energy onto the motion of *q_i_* and a measure of its cost. Kinetic energy flow is not used in the current analysis and is not discussed here [36].

To gain mechanistic insights, we need to look at how a mechanical quantity *A*(Γ) (e.g. potential energy flow) change as a reaction progresses. We first project *A*(Γ) onto a quantity *ξ*(Γ) that parameterizes the progress of a reaction, which we call the projector, then average over the ensemble of reactive trajectories:

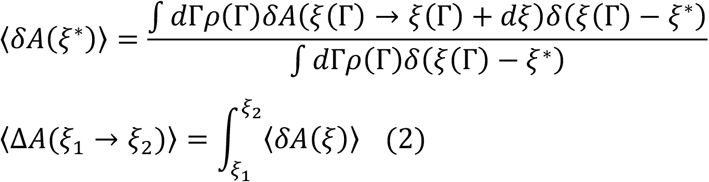

Here, 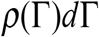 is the probability of finding the system in an infinitesimal volume *d*Γ around a phase-space point Γ in the reactive trajectory ensemble; *δ*(*x*) is the Dirac δ-function; *δA*(*ξ*(Γ) → *ξ*(Γ) + *dξ*) is the change in *A_i_* in a differential interval 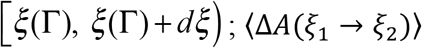 is the change in *A* in a finite interval [*ξ*_1_, *ξ*_2_]; Δ*A* can be Δ*W_i_* [29, 36]. The ensemble of reactive trajectories consists of trajectories that cover the transition period of a reaction process but exclude the waiting period [3].

### Generalized Work Functional

The source of complexity of protein dynamics is the highly entangled mechanical couplings between different coordinates. The potential energy flow through a coordinate *q_i_* provides an exact account of the energetic cost of its motion but does not inform us the effects of the mechanical couplings between different coordinates. These mechanical couplings are a consequence of the structure of Lagrange’s equation.

In generalized coordinates, the Lagrange’s equation is: 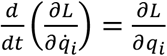, where L is the system Lagrangian *L* = *K* – *U*, defined as the difference between the kinetic energy 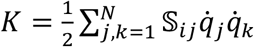 and the potential energy *U*. Here, 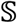 is the structural coupling tensor, defined as 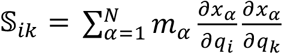, where the sum is over all the *N* coordinates in the system; *m_α_* and *x_α_* are the mass and a Cartesian coordinate of atom *α*. After expanding all the terms, we obtain:

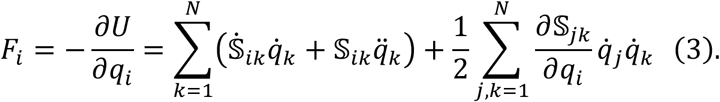

On the left-hand side is the force acting in the direction of 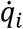 because 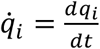 is in the same direction as the differential displacement *dq_i_*, which specifies the direction of 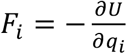 because *U* is a scalar. This force affects the motions of all the coordinates in the system due to the 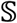 tensor. Without 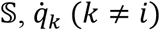 is orthogonal to 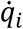 because generalized coordinates are orthogonal in any system configuration, but 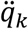 is not orthogonal to either 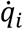 or 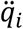 because coordinates are curvilinear in general. The 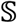 tensor rotates each 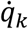 and 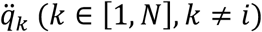 so that they all have overlaps with the direction of 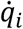 and *F_i_*, thus *F_i_* affects the motion of all the coordinates in the system. The effects of *F_i_* on *q_k_* and the effects of *F_k_* on *q_i_* are determined by the values of 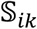 and 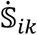, which are determined by the time-dependent system configuration.

Because *F_i_* affects motions of all coordinates, 〈*F_i_dq_k_*〉 (*i* ≠ *k*) provides an overall account of the impact of *F_i_* on the motion of *q_k_*, here 〈⋯〉 indicates average over the reactive trajectory ensemble. Accordingly, we define the generalized work flow from *q_i_* to *q_j_* as:

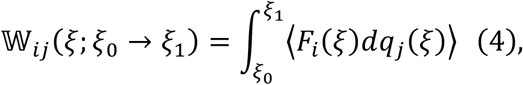

which is the aggregated impact of *F_i_* on the motion of *q_k_* as the reaction progresses from *ξ*_0_ to *ξ*_1_. The collection of all the 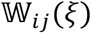 is a tensorial functional 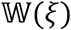 in the configuration space, which we call the generalized work functional because *F_i_dq_j_*(*i* ≠ *j*) is a generalization of the concept of mechanical work. For a specific value *ξ*_1_, 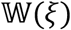 is a tensor 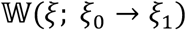, which we call the generalized work tensor.

With this definition, 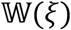 represents the integrated account of the effects of the pairwise mechanical couplings between different coordinates as a function of the progress of a reaction. Because 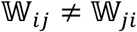, SVD is the proper tool for extracting the inherent structure of 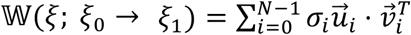, where *σ_i_* is the i-th singular value, 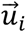 is the i-th left and 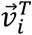 is the transpose of the i-th right singular vector. The leading term in this decomposition, 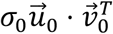, accounts for the dominant component of 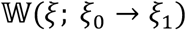. Accordingly, 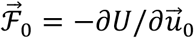 is the force in the direction of 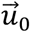. This force has the largest overall effects on the motions of different coordinates in the system. Because the 1D-RC is generally considered as the “driving force” of a reaction process [3], it is intuitively enticing to contemplate that it may coincide with 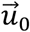. If this is the case, we have also found in GWF the physical determinant of the 1D-RC.

### An Example of Biomolecular Conformational Change

To test the effectiveness of the GWF in determining the 1D-RC, we applied it to the *C_7eq_* → *C_7ax_* isomerization of an alanine dipeptide in vacuum. This process is a prototype of conformational dynamics of proteins because alanine dipeptide is the smallest example of complex molecules. Here, we define a complex molecule as molecule whose non-reaction coordinates form a large enough heat bath for powering the RCs to cross the activation barrier [19, 24, 28, 29, 34, 37]. In contrasts, small molecules need an external energy source for activation, such as buffer gas in gas-phase and solvents in solution-phase reactions. The isomerization of an alanine dipeptide in vacuum carries some fundamental features that are unique to reactions in protein-like molecules but are absent in small molecules. Consequently, it has served as a standard system for testing methods for identifying RCs [3, 27, 28]. This isomerization process is mainly a rotation around the *ϕ* dihedral (Fig. 2). In previous studies, two backbone dihedrals *ϕ* and *θ*_1_ were identified as the RCs, as they can determine the committor value with satisfactory accuracy [28, 38].

**Figure 2:**
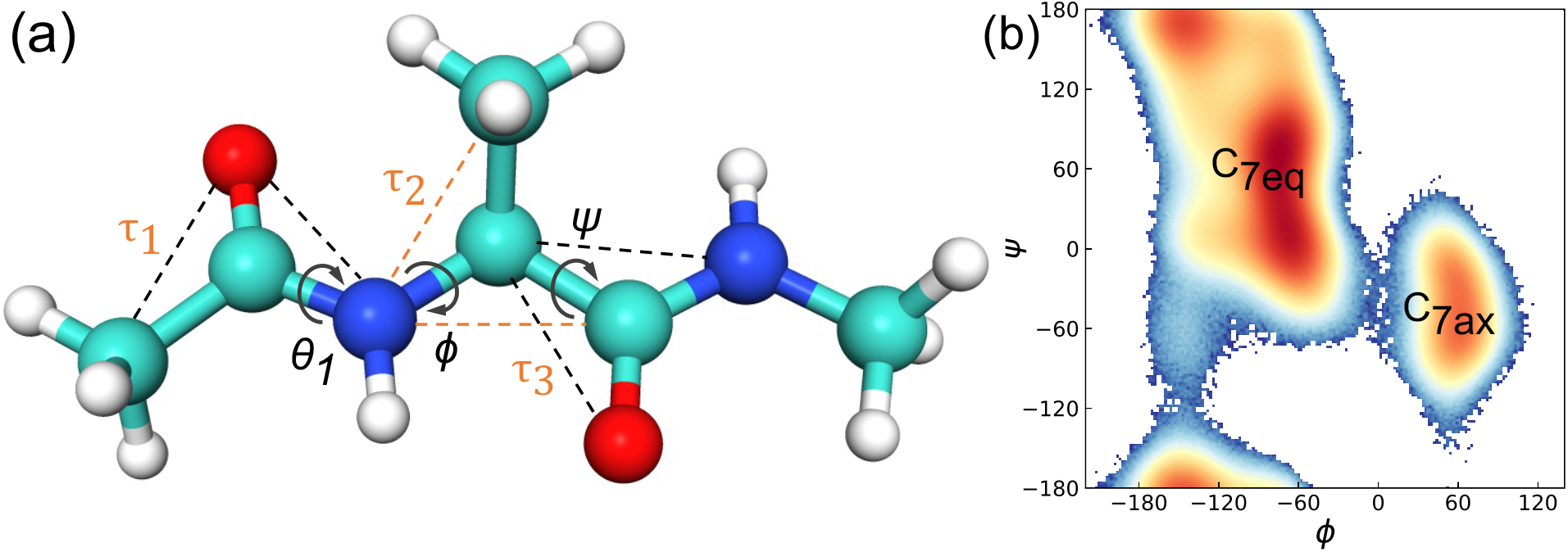
Schematics for the *C_7eq_* → *C_7ax_* isomerization of alanine dipeptide. **(a)** The molecular structure of alanine dipeptide. For a dihedral, its bond of rotation is marked by a curved arrow. For an improper dihedral, each of the two planes that define the it is spanned by 3 atoms, with the central atom chemically bonded to the other 2 atoms. We mark each plane by connecting the 2 atoms that are not bonded to each other with a dashed line. The edge shared by the 2 planes is the bond of rotation for the improper dihedral. **(b)** The definition of the *C_7eq_* and *C_7ax_* basins in the (*ϕ, ψ*)-plane. The heat map is the logarithm of the joint probability *p*(*ϕ, ψ*) obtained from a 6 *μs* equilibrium molecular dynamics simulation.

For analyses using energy flow and GWF, internal coordinates are the proper choice because they are the natural coordinates for describing protein motions and automatically satisfy all the restraints from the bonded interactions in a protein. In contrast, Cartesian coordinates cannot provide useful mechanistic information because their movements are dominated by strong restraint forces from bonded interactions that bear little relevance to the mechanism of protein dynamics.

### 1D-RC is Determined by the GWF

Figure 3a shows the generalized work functional for all the proper and improper dihedrals in the system computed with committor as the projector. The motion of the major RC, *ϕ*, is strongly affected by the forces from four coordinates: *ϕ* itself, the other known RC *θ*_1_, and two improper dihedrals *τ*_1_ and *τ*_2_ (Fig. 2a). Interestingly, both *τ*_1_ and *τ*_2_ are important players in transferring kinetic energy into *ϕ* based on our previous analysis of kinetic energy flows in this system [36]. In contrast, 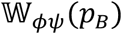 is significant but 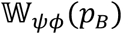 is small, suggesting that *F_ϕ_* has significant impact on the motion of *ψ* but *F_ψ_* has little influence on the motion of *ϕ*. This suggests that the motion of *ψ* is slaved to the motion of *ϕ* because the latter is the dominant factor for determining *F_ϕ_* [39, 40]. Consequently, *ψ*, though important for distinguishing the reactant and product states, is not an RC.

**Figure 3:**
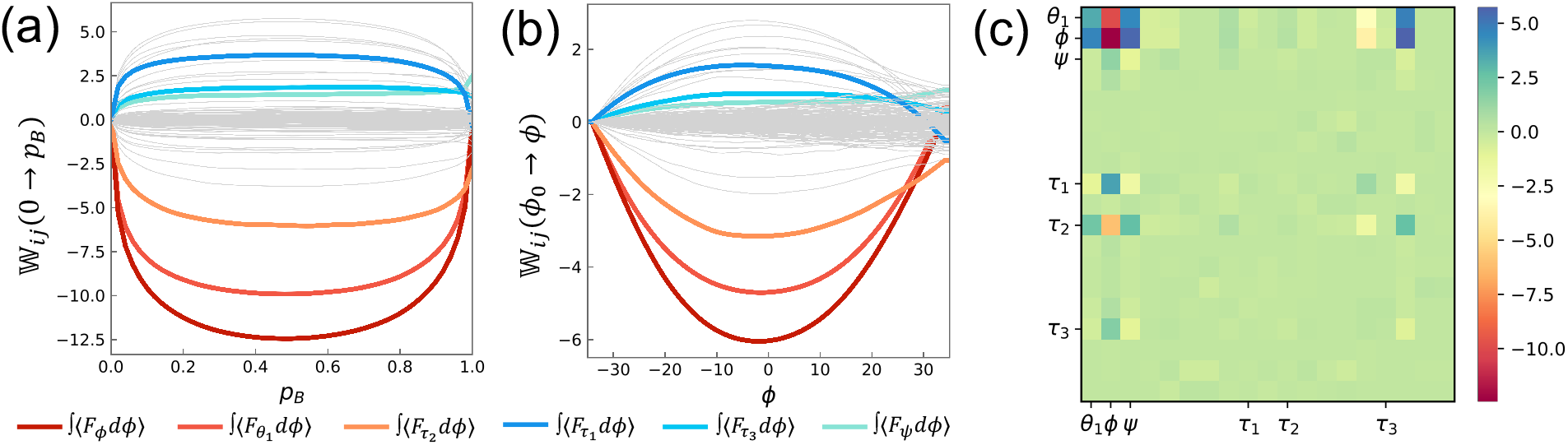
Generalized work flows and generalized work tensor. **(a)** Generalized work flows with *p_B_* as the projector for all the dihedrals and improper dihedrals in the system. There are 19 × 19 = 361 lines in the plot. The thick colored lines are 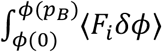 for *q_i_* that has significant magnitude, corresponding to the blocks with distinct colors in the second column of the generalized work tensor in panel (c). The rest are colored light gray. **(b)** Generalized work flows with *ϕ* as the projector. **(c)** The 19 × 19 generalized work tensor 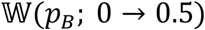 calculated for all the dihedrals and improper dihedrals in the system. Each tensor element is colored based on its value.

Figure 3c shows, 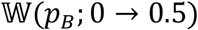, the generalized work tensor for *p_B_* = 0 to 0.5. We included all the dihedrals and improper dihedrals in constructing 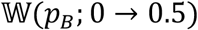, consisting of 19 coordinates in total. We did not include bonds or bond angles in 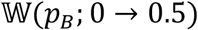 because the forces on them are of higher magnitude and oscillate very fast compared to forces on dihedrals and improper dihedrals. As a result, generalized work involving forces from bonds and bond angles have much higher noise.

The leading left singular vector of 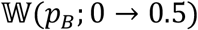 is 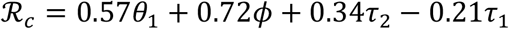. To test the quality of 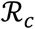, we carried out the standard “committor test” and compared the performance of 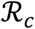 with RCs identified in previous studies [27, 28]. Because RCs must be sufficient for determining *p_B_*, configurations that share the same values of the RCs but differ in other coordinates should all have the same *p_B_* value. To test the quality of specific RCs, we harvest an ensemble of configurations that all have the values of the RCs corresponding to the transition state, with the other coordinates randomly distributed. The distribution of *p_B_* values for this ensemble should be peaked around 0.5 if the RCs can determine *p_B_*; a narrower distribution indicates higher quality of the RCs.

Figure 4a shows the results of the committor test. The orange line is the distribution of *p_B_* for configurations with *ϕ* = 0° and *ψ* = 40°, which corresponds to the value of *ϕ* at the transition state. This distribution peaks at *p_B_* = 0 and *p_B_* = 1, indicating that *ϕ* alone cannot determine committor [27, 28]. The green line is the distribution of *p_B_* for configurations with *ϕ* = 0° and *θ*_1_ = 3.6°. This combination was identified in previous studies as sufficient for determining committor. Indeed, this distribution peaks at *p_B_* = 0.5, though it is rather broad, indicating either missing RCs or the uncertainty inherent in the numerical calculation of *p_B_*. This is the limit reached by previous methods, both intuition-guided trial-and-error and machine-learning with neural network [27, 28].

**Figure 4:**
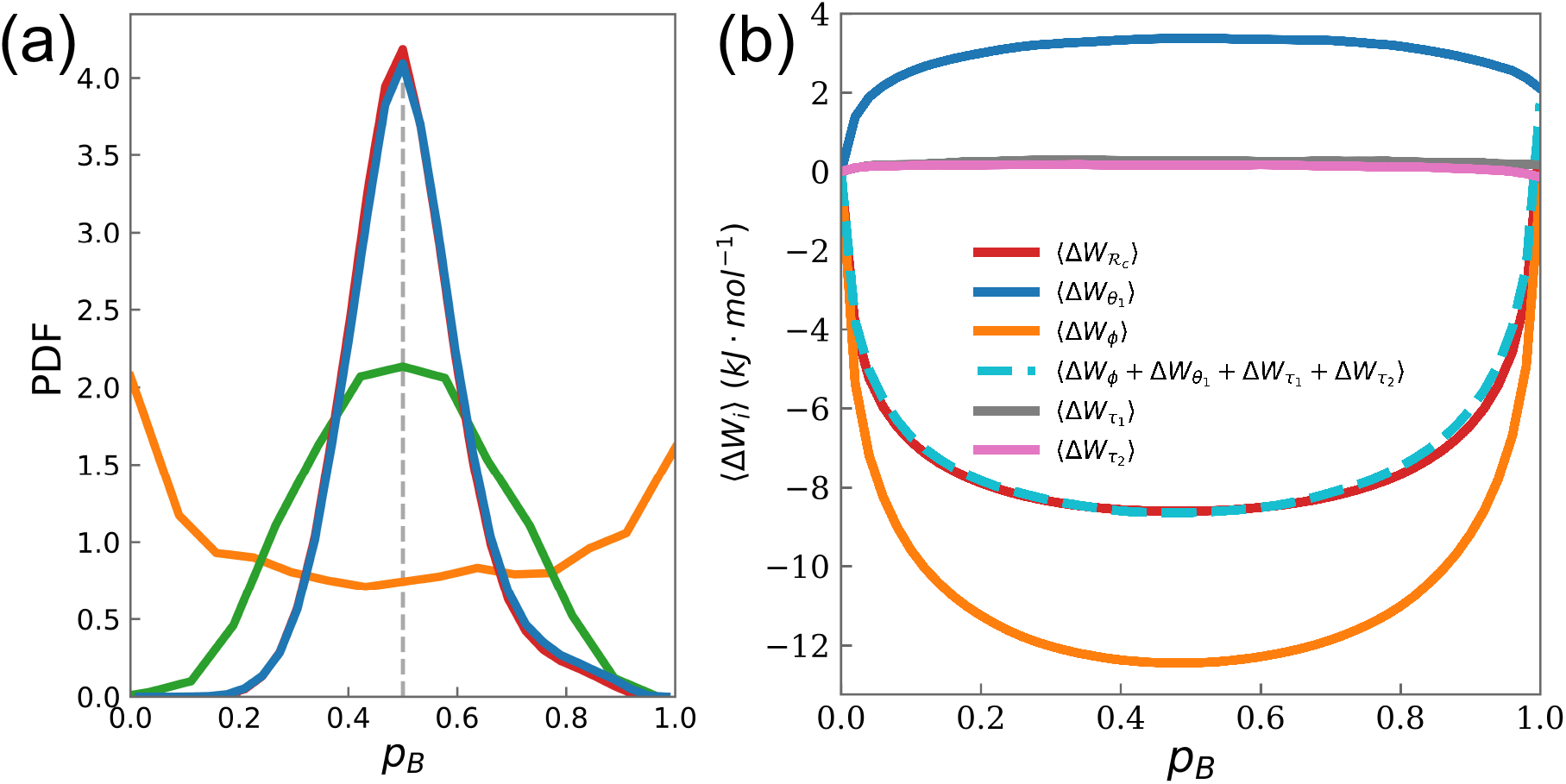
**(a)** Comparison of the results of committor tests for different RCs. **Orange**: PDF of *p_B_* values of the ensemble of configurations (*ϕ* = 0° and *ψ* = 40°) with *ϕ* as the RC, the same as used in ref. [27]. **Green**: PDF of *p_B_* values of the ensemble of configurations (*ϕ* = 0° and *θ*_1_ = 0.5°) with both *ϕ* and *θ*_1_ as the RCs, the same choice as in refs. [27, 28]. **Blue**: PDF of the ensemble of configurations with 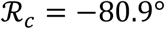. Red: PDF of the ensemble of configurations with 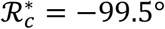. **(b)** Potential energy flows through *ϕ* (**orange**), *θ*_1_ (**blue**), *τ*_1_ (**pink**), *τ*_2_ (**grey**), and 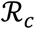 (**red**). The **cyan** dashed line represents Δ*W_ϕ_* + Δ*W*_*θ*_1__ + Δ*W*_*τ*_1__ + Δ*W*_*τ*_2__.

The blue line is the distribution of *p_B_* for configurations with 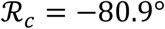. It sharply peaks at *p_B_* = 0.5 and has a narrow width, showing the high quality of 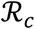. This significant improvement in the accuracy in determining committor value compared to previous results is due to two factors: 1) the inclusion of two improper dihedrals *τ*_1_ and *τ*_2_, 2) the correct coefficients in the analytical expression of 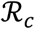. Because the effects of *τ*_1_ and *τ*_2_ on reaction dynamics are much weaker than those of *θ*_1_ and *ϕ*, which is shown by their smaller coefficients in 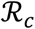, they are much more difficult to detect, explaining why they eluded both human intuition and machine learning [27, 28]. The fact that they are uncovered by 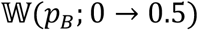 demonstrates the capability and precision of the GWF in capturing the essence of dynamics.

The red line is the distribution of *p_B_* for configurations with 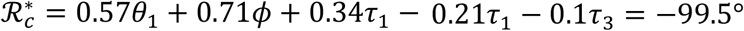, which almost identical to the blue line. This suggests that the improper dihedral *τ_3_* is a very minor RC, which is consistent with its small coefficient in 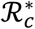.

Moreover, the matrix elements of 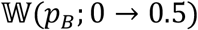 have inherent numerical uncertainty, especially that of *p_B_*. The fact 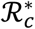 has minimal improvement over 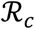 suggests that we have already reached the numerical limit, thus 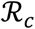 is likely the optimal 1D-RC. Since the GWF determines the rigorous 1D-RC, it is the correct physical operator for rigorous analysis of protein dynamics.

Figure 4b shows that 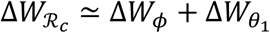. Because *ϕ* and *θ*_1_ carry the dominant majority of the total potential energy flow through the system based on previous studies [29, 36], this means 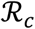 is responsible for the majority of the total potential energy flow through the system, confirming the 1D-RC as the single dominant channel for energy flows during a reaction.

### Is 1D-RC Curved?

An important conceptual question in reaction dynamics is whether the 1D-RC is straight or curved. There is no physical reason to expect that the 1D-RC should be straight, but in practice it is always assumed so for simplicity [24, 25]. This assumption is likely true for small molecules because the barrier top region is very narrow. For complex molecules, this assumption is less warranted due to the extended span of the barrier top. Since alanine dipeptide is the smallest complex molecule, it offers an excellent place to examine this matter.

Because the GWF in Fig. 3a is not linear, the direction of the 1D-RC constructed using 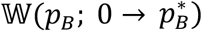 likely changes with the value of 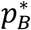, which means the direction of the 1D-RC in different intervals of committor value will be different. Figure 5a (solid black line) shows the projection of the concatenation of 1D-RCs for all the intervals of 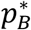 onto the (*ϕ*, *θ*_1_)-plane, which amounts to a piecewise linearization of the true 1D-RC. We choose this projection because it is easy to visualize in a plane, and *ϕ* and *θ*_1_ are the dominant RCs. Indeed, the 1D-RC is slightly curved. This curvilinearity is quantified in Fig. 5b (orange line), which shows the slope of the 1D-RC as a function of *p_B_*. Figure 5c shows the results of the committor test on the curved 1D-RC. Indeed, it can predict committor value with high accuracy over the entire range of *p_B_*.

**Figure 5:**
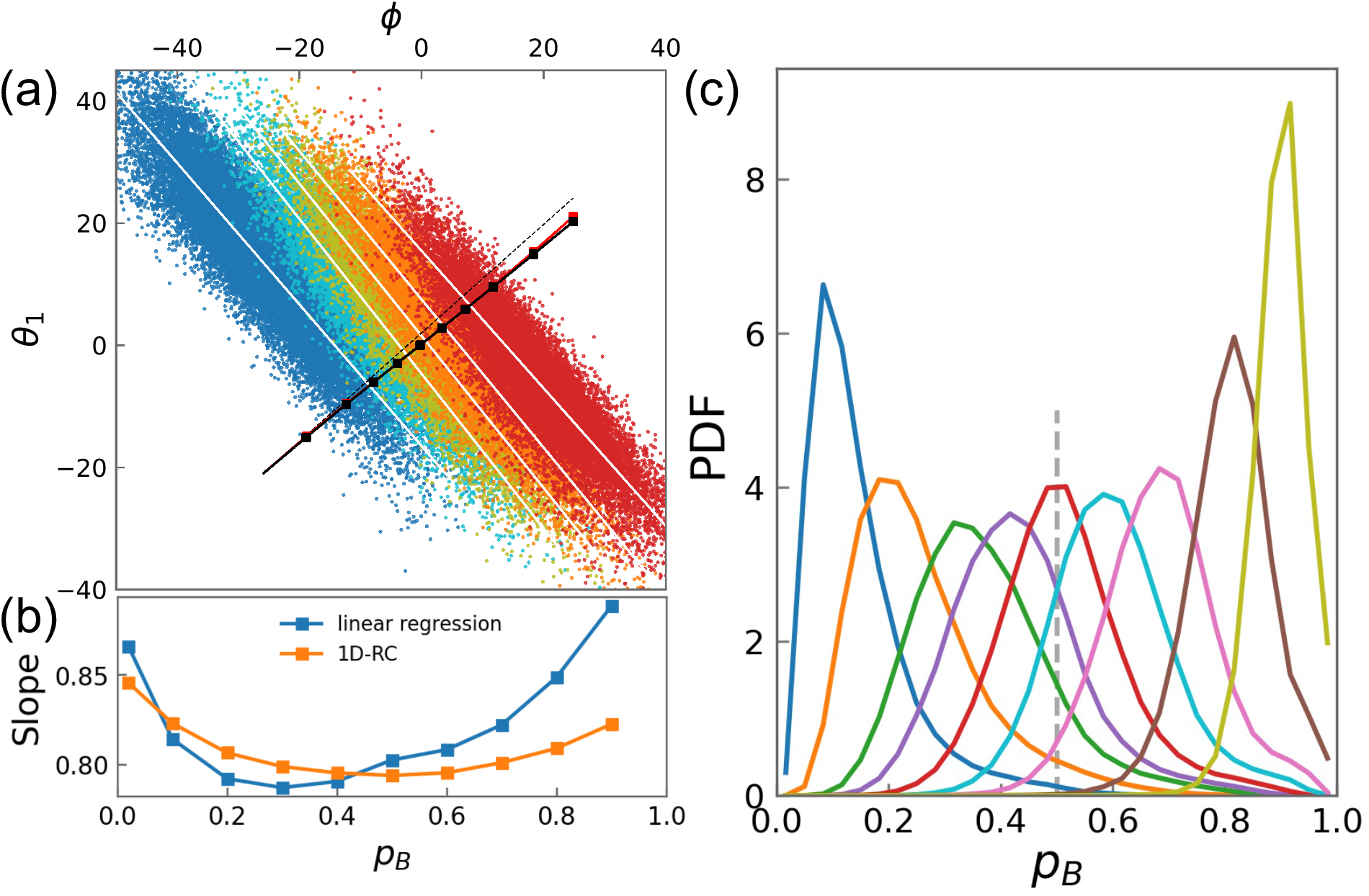
The curvedness of 1D-RC. **(a)** The scattered points represent projection of ensembles of configurations with *p_B_* = 0.02,0.3,0.5,0.7,0.9 onto the (*ϕ*, *θ*_1_)-plane. The white straight line through a colored “cloud” is its linear regressions, thus each one represents an approximation to the intersection of the corresponding iso-committor surface with the (*ϕ*, *θ*_1_)-plane. The solid black line is 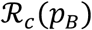 constructed as a concatenation of 10 short line segments; each segment is the 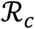 for an interval of *p_B_*. The 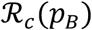 for a specific 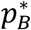 is the leading left singular vector of the tensor 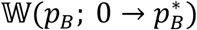. To construct the short line segments corresponding to different values of *p_B_*, we first prepare 11 ensembles of configurations; each one contains configurations of a specific *p_B_* value: *p_B_* = 0.02, 0.1, 0.2, 0.3, 0.4, 0.5, 0.6, 0.7, 0.8, 0.9, 0.98. Each ensemble of configurations are then projected onto the (*ϕ*, *θ*_1_)-plane and the center of mass, *c_i_* (*i* = 1, …, 11), of each ensemble is computed. The *ϕ* coordinate of this series of *c_i_* defines a sequence of positions along the *ϕ*-direction: *ϕ_i_* =-18.9°, −12.3°, −7.8°, −3.9°, −0.2°, 3.3°, 7.2°, 11.7°, 18.4°, 24.8°. We start from *c*_1_ as the starting point of the first line segment, with its slope as the slope of 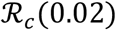 projected onto (*ϕ*, *θ*_1_)-plane. The first segment ends at 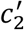, with its *ϕ*-coordinate equal to *ϕ*_2_, and its *θ*_1_-coordinate equal to: 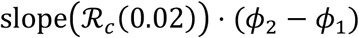. The other segments are constructed in a similar way. The red solid line is constructed in the same manner as the black solid line, but the direction for each line segment is chosen as the direction normal to the linear regression of each *p_B_* ensemble. The black dashed line is a straight line in the direction normal to the linear regression of the ensemble of *p_B_* = 0.9. Both solid lines deviate from this straight dashed line, demonstrating the curvedness of the 1D-RC. The black and red solid lines are essentially coincidental, showing that 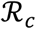 is perpendicular to the isocommittor surfaces throughout the transition region. **(b)** The slopes of the line segments in panel (a) as function of committor. **(c)** Results of committor tests for ensembles of configurations with 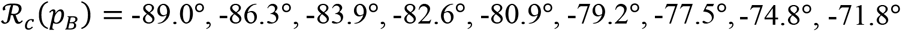, corresponding to *p_B_* = 0.1, 0.2, 0.3, 0.4, 0.5, 0.6, 0.7, 0.8, 0.9, respectively.

To better understand this curvilinearity of the 1D-RC, we projected five ensembles of configurations, each one with *p_B_* in the range of 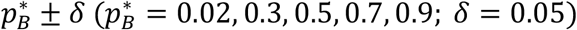, onto the (*ϕ*, *θ*_1_)-plane. Each ensemble consists of configurations on an iso-committor surface, with the thickness of the surface defined by *δ*. It is often assumed that an iso-committor surface is a hyperplane in the configuration space [24, 25, 32]. If this is true, each ensemble of configurations should form a straight strip with a width determined by *δ* in the (*ϕ*, *θ*_1_)-plane. Instead, each ensemble of configurations clusters into an ellipse. To find out the intersections of each iso-committor surface with the (*ϕ*, *θ*_1_)-plane, we obtain a straight line through each ellipse by linear regression. Figure 5a (white lines) shows that the direction of these iso-committor lines changes slowly with 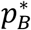. The red solid curve is the concatenation of the normal directions of all the iso-committor lines; it is slightly bent. Intriguingly, it essentially coincides with the black curve, suggesting that the 1D-RC, though slight curved, is always perpendicular to the sio-committor surface.

### Computation of the GWF is Robust and Cost-Effective

In the results above, the GWF was computed using committor as the projector. Since numerical evaluation of *p_B_* is computationally expensive [41], it could limit the utility of the current method as a practical approach for identifying the 1D-RC in large systems. To overcome this potential limitation, we computed the GWF using *ϕ* as the projector, which is a good order parameter for distinguishing the reactant and product basins, but insufficient for parameterizing the progression of the reaction (Fig. 4a). Since RCs control the dynamics of the transition period, not the dynamics within the reactant basin, we only consider the range of *ϕ* ∈ [-35°, 35°], corresponding to the region with *p_B_* ∈ (0,1).

Figure 3b shows 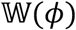, the GWF calculated with *ϕ* as the projector, which differ from 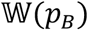 in Fig. 3a. This is expected because projector has a significant impact on the ensemble average and the relation between *ϕ* and *p_B_* is neither linear nor one-to-one, the latter because *ϕ* is not sufficient for determining *p_B_*. Importantly, the leading left singular vector of the generalized work tensor 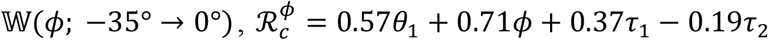, is essentially identical to 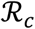, thus it can determine *p_B_* with the same accuracy. Therefore, our method can be used to identify the correct 1D-RC using an order parameter as the projector for computing the GWF. This tremendously decreases the computational cost of the current method compared to previous methods, as now the major computational cost is to harvest sufficient number of reactive trajectories so that the computation of the GWF will converge. More importantly, substituting *p_B_*, which accurately parameterizes the progress of a reaction but is computationally expensive, with an order parameter, which only distinguishes stable basins but is computationally very efficient, has little impact on the accuracy for identifying the 1D-RC. This fact makes the current approach a robust yet cost-effective method for identifying the 1D-RC in complex molecules.

### Origin of Collectivity in Functional Protein Dynamics

Aside from providing a practical method for identifying 1D-RC, the GWF also answers a fundamental question in protein dynamics. Functional protein dynamics often involve significant changes in the global protein structure, which are generally assumed to be a result of collective motions. This type of collective motion is different from normal modes, which can account for small amplitude vibrations around equilibrium structures but cannot produce conformational changes. On the other hand, normal modes have a clear physical origin as the eigenvectors of the Hessian. In contrast, it remains a puzzle what the collective modes responsible for protein conformational changes are and what their physical origin is.

Since the 1D-RC fully determines the progress of functional protein dynamics, by definition it is the collective mode responsible for the conformational change of a protein. Each internal coordinate provides the natural description of a local movement of a protein and aggregation of many local movements generate the global conformational change. These local movements are mechanically coupled together by the 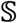 tensor, synergizing them with each other. This is the reason behind the collectivity in functional protein dynamics. The gist of the time-dependent mechanical couplings introduced by the 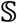 tensor is accurately encapsulated by the GWF, which has a simple mathematical structure—an asymmetric tensor. The left singular vectors of the generalized work tensor are the directions of forces that have persistent effects on conformational dynamics, as they survive the ensemble average in Eq. (4). The leading singular vector is the direction with the most significant effects; thus it is the most important collective mode for the conformational change. The major components of this leading singular vector and the synergy among these components, defined by their coefficients in 1D-RC, are the precise definition of the collectivity responsible for functional dynamics. This is the physical reason that protein conformational change is achieved through a global collective mode. Because the GWF determines the 1D-RC, it is also the physical origin of collectivity in functional protein dynamics.

### A Tentative Roadmap for the Structure-Function Relationship of Proteins

The critical link between the structure and function of a protein is its functional dynamics. We uncovered the GWF as the physical determinant of the 1D-RC that controls functional dynamics. The origin of GWF is that the force on one coordinate affects the motions of all the coordinates in the protein due to the 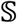 tensor. Both the forces and the 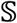 tensor of a protein are determined by its structure; thus we obtain a chain of command that connects the structure of a protein to its function.

As a result, we have a tentative roadmap for the structure-function relationship (Fig. 1b). The structure of a protein determines both atom-atom interactions and the 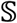 tensor. Together, they determine the GWF, which determines the dominant forces in the protein and the RCs. The RCs determine the functional dynamics, thus determines the function.

## Discussions

In this paper, we developed a rigorous and computationally efficient method for identifying the rigorous 1D-RC. The 1D-RC for the *C_7eq_* → *C_7ax_* isomerization of an alanine dipeptide in vacuum is a linear combination of two dihedrals, *ϕ* and *θ*_1_, that were known from previous studies, and two improper dihedrals *τ*_1_ and *τ*_2_, that are identified for the first time. The 1D-RC can predict the committor value over the entire transition region (*p_B_* ∈ (0,1)) with an accuracy far exceeding what was achieved before, attesting the value of rigorous physics-based method. In addition, the new method allows us to determine the 1D-RC as a function of *p_B_*, revealing that the 1D-RC slightly curved instead of being straight.

Most importantly, the 1D-RC is determined by a simple and general mechanical operator, the generalized work functional. Since the progress of a reaction in a protein is controlled by a 1D-RC, it means the reaction dynamics has an intrinsic order underneath its apparent chaos and complexity. The GWF is the determinant of this intrinsic order because determines the 1D-RC, which encapsulates this intrinsic order. The complexity of protein dynamics originates from complexity of the couplings between different coordinates. The underlying operator for this coupling is the structural coupling tensor 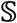, which makes force of one coordinate directly affect the dynamics of many other coordinates by rotating their velocity and acceleration vectors. The mechanical consequences of these couplings are encapsulated in the GWF. Singular value decomposition on the generalized work tensor unveils its inherent structure, thus reveals the underlying order of protein dynamics. As a result, the GWF is also the physical origin of collectivity in protein dynamics and enabled us to build a tentative roadmap for the structure-function relationship of proteins.

Finally, RCs play a central role in the efficiency of algorithms for enhanced sampling of the configuration and trajectory space of proteins. Because our method enables identification of rigorous 1D-RC with modest computational cost, it can potentially point to new directions for developing algorithms for enhanced sampling.

## Methods

All simulations were performed using the molecular dynamics software suite GROMACS ^[42]^ with transition path sampling implemented. Amber 94 force field was used to facilitate comparison with previous results [29, 36, 38, 43, 44]. The structure of the alanine dipeptide was minimized using steepest descent algorithm and heated to 300 K using velocity rescaling with a coupling constant of 0.2 ps. The system was then equilibrated for 200 ps and no constraints were applied. The time step of integration was 1 fs. Basin *C_7eq_* is defined as −190° < *ϕ* < −55° and −60° < *ψ* < 190°; basin *C_7ax_* is defined as 50° < *ϕ* < 100° and −80° < *ψ* < 0°. We used transition path sampling method to generate the ensemble of reactive trajectories between these two basins that are used in all the analyses discussed here [37]. All the averaged quantities discussed in the text were averaged over 20,000 trajectories.

